# Human iPSC-derived muscle cells as a new model for investigation of EDMD1 pathogenesis

**DOI:** 10.1101/2024.12.08.627381

**Authors:** Marta Lisowska, Marta Rowińska, Aleksandra Suszyńska, Claudia Bearzi, Izabela Łaczmańska, Julia Hanusek, Amanda Kunik, Volha Dzianisava, Ryszard Rzepecki, Magdalena Machowska, Katarzyna Piekarowicz

## Abstract

Emery-Dreifuss muscular dystrophy type 1 (EDMD1) is a rare genetic disease caused by mutations in the *EMD* gene, which encodes the nuclear envelope protein emerin. Despite understanding the genetic basis of the disease, the molecular mechanism underlying muscle and cardiac pathogenesis remains elusive. Progress is restricted by the limited availability of patient-derived samples, therefore there is an urgent need for human-specific cellular models.

In this study, we present the generation and characterization of induced pluripotent stem cell (iPSC) lines derived from EDMD1 patients carrying *EMD* mutations that lead to truncated or absent emerin, together with iPSCs from healthy donor. The patient-specific iPSCs exhibit stable karyotypes, maintain appropriate morphology, express pluripotency markers and demonstrate the ability to differentiate into three germ layers.

To model EDMD1, these iPSCs were differentiated into myogenic progenitors, myoblasts and multinucleated myotubes, which represent all stages of myogenesis. Each developmental stage was validated by the presence of stage-specific markers, ensuring the accuracy of the model. We present the first iPSC-based *in vitro* platform that captures the complexity of EDMD1 pathogenesis during myogenesis. This model can significantly contribute to understanding disease mechanisms and develop the targeted therapeutic strategies for EDMD1.

## 1. Introduction

Emery-Dreifuss muscular dystrophy type 1 (EDMD1, OMIM #310300) is a rare genetic disorder with a prevalence of 1 per 100,000 male births. It is an X-linked recessive disease caused by mutations in the *EMD* gene (*locus Xq28*) coding for nuclear envelope (NE) protein emerin [1]. The main symptoms of EDMD1 are observed in the skeletal muscles and heart, including muscle weakness and wasting, tendon contractures, and cardiac dysfunction [2]. The disease is incurable, with a substantial risk of sudden death in middle age caused by heart block. It is still unclear whether the changes observed in progenitor cells or at the later stages of myogenesis are crucial for aberrations in muscle development and regeneration in EDMD1. Understanding the disease mechanisms is fundamental for design, development, and evaluation of the therapy.

Emerin is a transmembrane protein with a LAP2, emerin, MAN1 (LEM) domain that interacts with various partners in the cell nucleus, including the highly conserved DNA-binding protein BAF (Barrier-to-Autointegration Factor), facilitating DNA tethering to NE. Emerin plays an important role in nuclear stability, gene regulation, chromatin organization, and the cell cycle [3–5]. It is especially important in striated muscle cells, where emerin is essential for muscle development, maintenance and regeneration [6–8]. The absence or misexpression of emerin leads to muscle degeneration. Furthermore, emerin, together with other NE protein, lamin A/C, appears critical for muscle cells differentiation [9]. Dysfunction of satellite cells (SCs), the progenitors of muscle cells, has also been hypothesized to contribute significantly to the progression of EDMD by impairing myofiber repair and regeneration [10]. Despite this, the role of emerin in precursor muscle cells remains poorly understood. One major limitation is the lack of appropriate models for studying emerin role in pathogenesis. Not enough data has been acquired with human, patient-derived muscle precursor cells, with an investigation of patients-derived SCs mostly restricted to microscopic analysis of rare muscle biopsies [11–13].

The function and regulation of emerin are species-dependent, limiting the utility of mouse models and non-human cells [14–15]. While emerin-null mice exhibit only minor phenotypic changes, such as delayed skeletal muscle regeneration and repair, mild atrioventricular alterations, and motor coordination defects, humans with emerin deficiency develop muscular dystrophy. Additionally, mutation-related phenotype in humans significantly depends on genetic background [16–18]. It is possible to preserve it using an approach based on MyoD overexpression in patient’s fibroblasts to obtaining myoblasts [19], but this model does not allow for studying myogenic precursors, as well as MyoD-related signalling pathways may be influenced. The absence of accessible human-derived muscle tissue is a significant barrier for investigating EDMD1 *in vitro*. The availability of patients’ skeletal muscle biopsies is very limited, and especially patients’ cardiomyocytes for research are not available. Therefore, patient-specific induced pluripotent stem cells (iPSCs) offer a promising alternative for providing human muscular *in vitro* models. Importantly, as the starting material is derived from patients, the model retains the patient-specific genetic background, providing a physiologically relevant system. Differentiated muscle cells derived from iPSCs could serve as a key model for EDMD1, to elucidate disease mechanisms and the development of targeted therapies.

iPSCs can be differentiated into striated muscle progenitors and subsequently into myoblasts and multinucleated myotubes using a transgene-free protocol [20]. This approach enables the generation of sufficient amount of research material. The initial step of iPSCs differentiation procedure produces muscle progenitors, also called SCs. However, this method does not allow for obtaining separately quiescent and activated SCs populations. Single-cell transcriptomic analyses of muscle progenitors obtained *via* iPSCs differentiation using various chemical methods revealed that their expression profiles closely resemble those of embryonic skeletal muscle progenitor cells (SMPCs) while also sharing substantial similarity with adult muscle SCs population [21].

SCs are typically characterized by the expression of the transcription factors belonging to the paired box family – Pax3 and Pax7, which serve as key markers of this cell population. They are both essential regulators of myogenesis. Pax3 is involved in the early stages of muscle development by regulating downstream regulatory factors MyoD and Myf5. Pax7 is dominant in postnatal skeletal muscle differentiation. It is responsible for regulating expansion, maintenance and proliferation of SCs. It is also involved in maintaining SCs in an undifferentiated state. The combined action of Pax3 and Pax7 optimizes the conversion of stem cells into muscle cells [22–24].

iPSC-derived myogenic progenitors can be further differentiated to myoblasts, followed by differentiation to myocytes and the formation of multinucleated myotubes, which serve as an *in vitro* model for mature skeletal muscle cells. The differentiation process is closely associated with changes in transcriptional profiles. Myoblasts are usually characterized by the expression of transcription factors MyoD and Myf5, while typical markers of myotubes are myosin heavy chain (MHC), actinin, titin or elevated desmin level.

In this study, we generated and characterized five new iPSC lines from fibroblasts obtained from EDMD1 patients and a healthy donor. These included one clone from a patient bearing an *EMD* mutation c.153del and two clones from a patient with mutation c.451dup, both causing a frameshift and creation of a premature termination codon, resulting in the lack of protein or shorter protein, respectively. Subsequently, we differentiated the newly established clones, along with four iPSC clones previously generated by reprogramming fibroblasts from two EDMD1 patients with c.153del mutation [25], into skeletal muscle cells. Using a transgene-free differentiation method, we successfully obtained cells representing three stages of skeletal muscle development: myogenic progenitors, myoblasts and myotubes. The differentiated muscle cells expressed proper stage-specific markers. Additionally, we obtained myotubes, the final stage of *in vitro* muscle differentiation, showing sarcomere structure and multiple nuclei per cell, which are the features of mature skeletal muscle cells. We also observed the delayed differentiation of emerin-null cells, as those myoblasts needed more time to be able to create myotubes.

This work establishes a novel, robust model of EDMD1, providing a valuable platform for investigating disease pathogenesis and developing targeted therapies. Furthermore, the generated iPSCs may also be used in the future for the generation of cardiac and nerve cells, and development of multilineage organoids. The platform could be extent by preparation of isogenic controls to take into account the impact of genetic background on disease development.

## 2. Results

For iPSCs generation, we used dermal fibroblasts obtained from a skin biopsy of two EDMD1 patients and a healthy donor. Donors were male, Caucasian ethnicity, and unrelated. The patients had confirmed mutation in *EMD* gene (*locus Xq28*) and exhibited typical EDMD symptoms [26–28].

We reprogrammed the fibroblasts to iPSCs using non-integrating Sendai virus (SeV) vectors encoding the four Yamanaka factors: Oct3/4, Sox2, Klf4, and c-Myc. We established two iPSC clones for the healthy donor, two clones for the patient with *EMD* mutation c.451dup, and one clone for the patient with *EMD* mutation 153del (Table 1). Additionally, we previously generated four clones from two patients with *EMD* mutation c.153del [25], which were also used in this study (Table 1).

**Table 1.** Summary of generated iPSC clones. The Table summarizes newly validated iPSC clones from three donors alongside previously reported clones from two donors. Emerin protein level was evaluated by WB analysis in myotubes (Figure 4C).

| iPSCs line name | Unique stem cell line identifier | Disease | Age of biopsy | Genotype (*EMD* gene) | Emerin expression |
| --- | --- | --- | --- | --- | --- |
| Newly established clones |  |  |  |  |  |
| C1M35 L26 | UWRBTi001-C | N/A | 35 | NM_000117.3(EMD), CCCCCCAGC, GGGAACGCC | normal |
| C1M35 1.6 | UWRBTi001-D | N/A |  |  |  |
| E1M40 1.4 | UWRBTi003-A | EDMD1 | 40 | NM_000117.3(EMD):c.153del, p.(Ser52fs), CCCCC_AGCT, hemizygous, rs876661345 | not detected |
| E1M19 1.5 | UWRBTi005-A | EDMD1 | 19 | NM_000117.3(EMD):c.451dup*, p.(Glu151fs), GGGGAACGCC, hemizygous | truncated protein, lower level |
| E1M19 1.8 | UWRBTi005-B | EDMD1 |  |  |  |
| Previously generated clones |  |  |  |  |  |
| E1M40 1.7 | UWRBTi003-B | EDMD1 | 40 | NM_000117.3(EMD):c.153del, p.(Ser52fs), CCCCC_AGCT, hemizygous, rs876661345 | not detected |
| E1M40 1.9 | UWRBTi003-C | EDMD1 |  |  |  |
| E1M51 1.4 | UWRBTi004-A | EDMD1 | 51 | NM_000117.3(EMD):c.153del, p.(Ser52fs), CCCCC_AGCT, hemizygous, rs876661345 | not detected |
| E1M51 1.8 | UWRBTi004-B | EDMD1 |  |  |  |

\*Mutation c.451dup was previously reported also as c.450insG or c.450dup (e.g. www.umd.be/emd database).

iPSCs were expanded to at least passage 10, characterized and validated, as summarized in Table 2. Other iPSC clones were discarded due to detection of genomic instability. The morphology of colonies was typical for iPSCs [29–31]. The iPSC colonies had well-defined edges. They were bright and shiny under phase-contrast microscopy. Cells were round, tightly packed and had large nuclei (Figure 1A). We confirmed the expression of pluripotency markers using immunofluorescence (IF) staining (Figure 1B; control negative staining shown in Supplementary Figure S1) and quantitative PCR (qPCR, Figure 1C). All clones expressed *OCT4, SSEA4, NANOG, hTERT, LIN28, SOX2* and *REX-1*.

**Table 2.**
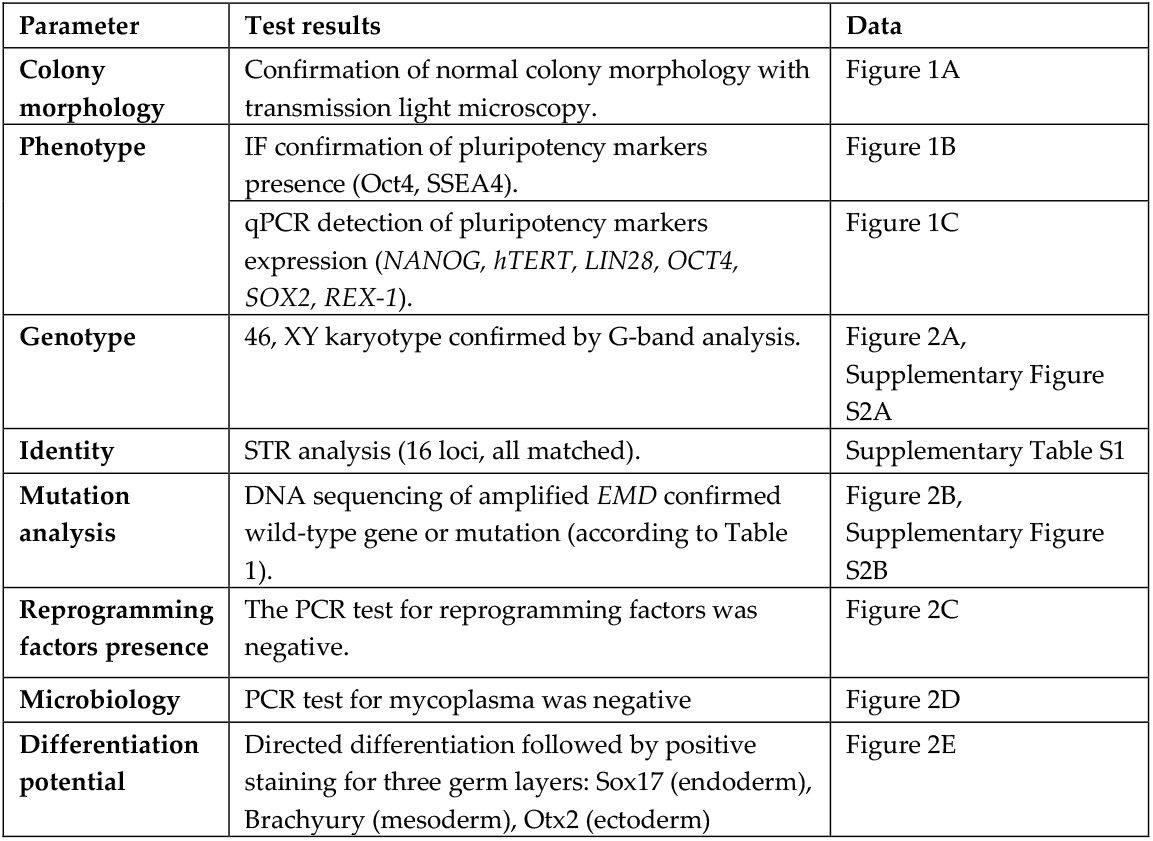
Characterization and validation of iPSCs. The Table contains all tests performed for iPSCs validation with reference to corresponding data.

| Parameter | Test results | Data |
| --- | --- | --- |
| **Colony morphology** | Confirmation of normal colony morphology with transmission light microscopy. | Figure 1A |
| **Phenotype** | IF confirmation of pluripotency markers presence (Oct4, SSEA4). | Figure 1B |
|  | qPCR detection of pluripotency markers expression (*NANOG*, *hTERT*, *LIN28*, *OCT4*, *SOX2*, *REX-1*). | Figure 1C |
| **Genotype** | 46, XY karyotype confirmed by G-band analysis. | Figure 2A, Supplementary Figure S2A |
| **Identity** | STR analysis (16 loci, all matched). | Supplementary Table S1 |
| **Mutation analysis** | DNA sequencing of amplified *EMD* confirmed wild-type gene or mutation (according to Table 1). | Figure 2B, Supplementary Figure S2B |
| **Reprogramming factors presence** | The PCR test for reprogramming factors was negative. | Figure 2C |
| **Microbiology** | PCR test for mycoplasma was negative | Figure 2D |
| **Differentiation potential** | Directed differentiation followed by positive staining for three germ layers: Sox17 (endoderm), Brachyury (mesoderm), Otx2 (ectoderm) | Figure 2E |

**Figure 1.**
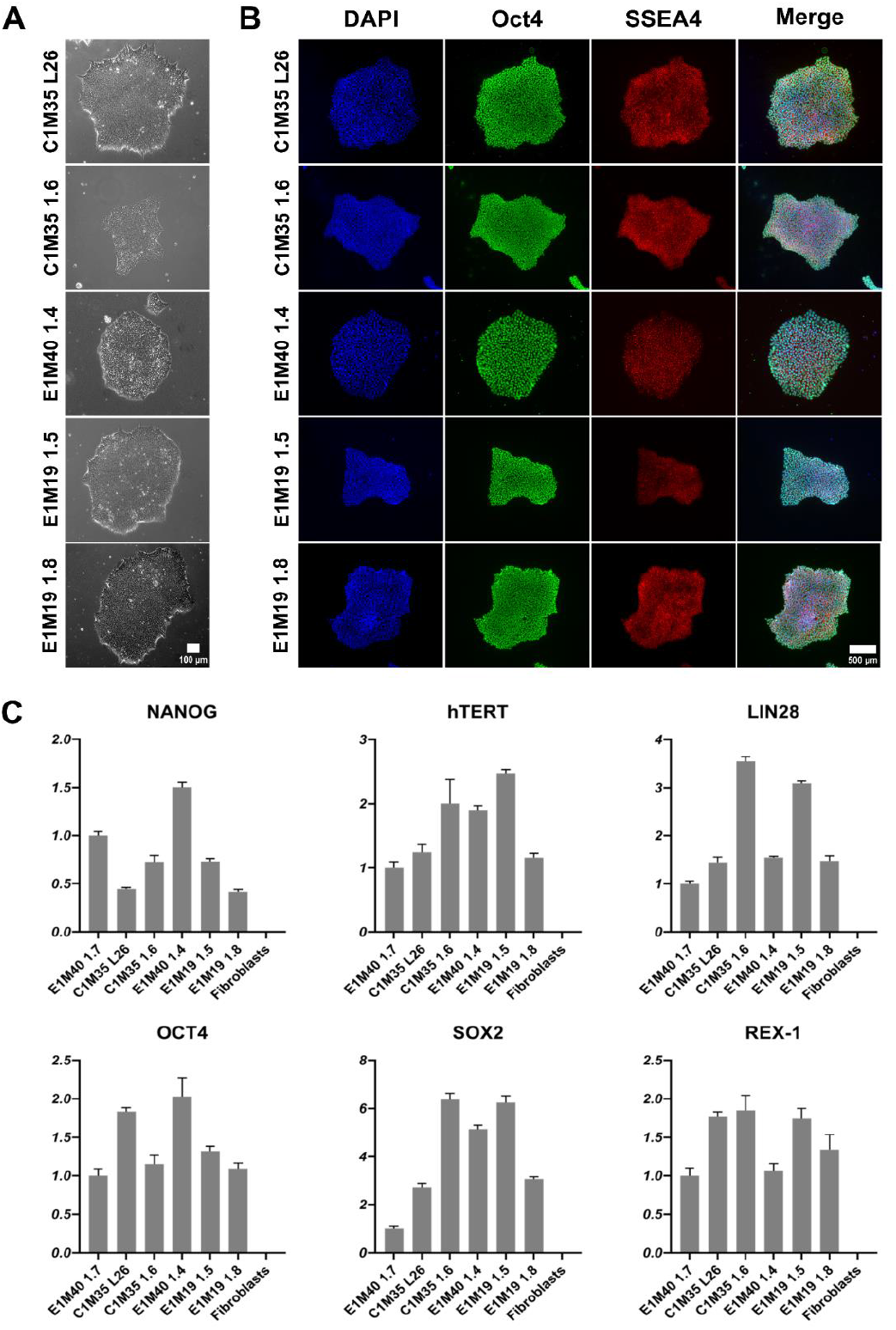
Morphology and pluripotency markers expression analysis of iPSC lines. **(A)** Images show the morphology of properly formed iPSC colonies in transmitted light (10x objective). The scale bar is 100 μm. **(B)** IF staining of iPSCs shows proper localization and equal distribution of pluripotency markers Oct4 (AF488) in nucleus and SSEA4 (TRITC) in cell surface of representative iPSC colonies. Control negative staining of HeLa cells is shown in Supplementary Figure S1. The scale bar is 500 μm. **(C)** qPCR analysis shows relative gene expression levels of iPSC pluripotency markers, normalized to the previously published E1M40 1.7 clone [25] and to glyceraldehyde-3-phosphate dehydrogenase (*GAPDH)* expression level. Error bars present standard deviations, n=4 (technical replicates).

The genomic stability of iPSC clones was confirmed by karyotypes analysis (Figure 2A) and short tandem repeats (STR) profiling (16 loci, Supplementary Table S1), compared to parental fibroblasts (Supplementary Figure S2A). DNA sequencing of the *EMD* gene confirmed the wild-type sequence in control cells and the *EMD* mutations in EDMD1 cells (Figure 2B), matching the sequences found in the parental fibroblasts (Supplementary Figure S2B).

**Figure 2.**
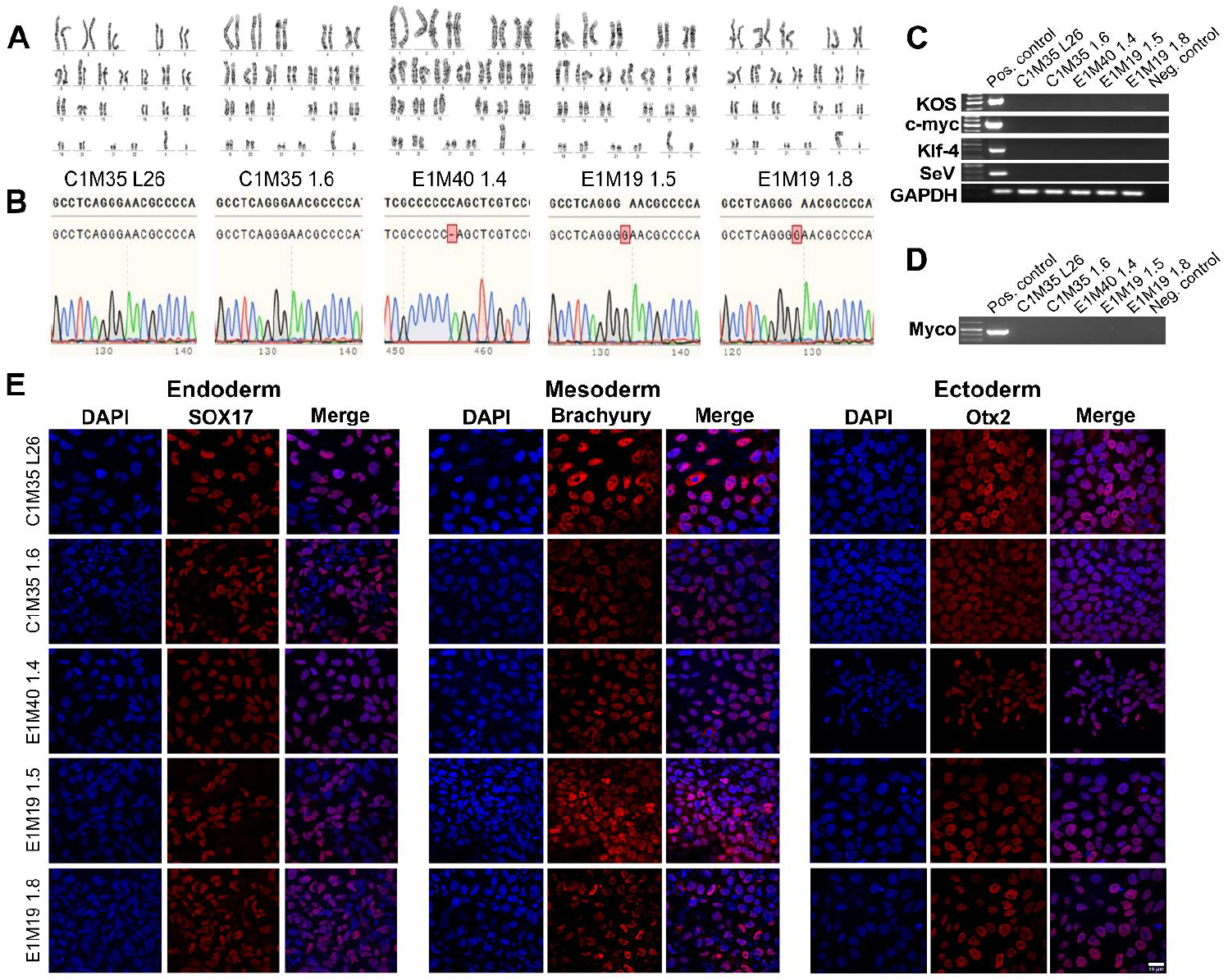
Validation of the iPSC lines – genetics, purity and differentiation ability. **(A)** G-banding analysis shows proper 46, XY karyotypes. **(B)** DNA sequencing chromatogram of fragment of amplified *EMD* gene confirmed wild-type sequence in control lines, cytidine deletion c.153del (CCCCC_AG) in E1M40 1.4 clone and guanine duplication c.451dup (GGG**G**AAC) in both E1M19 lines. The upper sequence shows the reference wild-type *EMD*, while the bottom sequence shows the results of DNA sequencing. **(C)** PCR verification shows the loss of each reprogramming vector (KOS, c-MYC, KLF4, SeV) in iPSC lines, together with positive control GAPDH. **(D)** PCR analysis shows lack of mycoplasma contamination in each iPSC clone. **(E)** IF staining shows germ layers markers. Each iPSC clone was differentiated independently into three germ layers on coverslips, followed by cells staining for adequate markers: Sox17 (Cy5) for endoderm, Brachyury (Cy5) for mesoderm and Otx2 (Cy5) for ectoderm. Control negative staining of undifferentiated iPSCs is shown in Supplementary Figure S3. The scale bar is 20 μm.

The absence of SeV vectors used for reprogramming was confirmed around passage 15 by PCR detecting four different sequences of vectors (Figure 2C). All cells were confirmed by PCR as free of mycoplasma (Figure 2D). Finally, we checked the ability of our clones to differentiate *in vitro* into three germ layers, using dedicated growth media (as described in the *Materials and Methods* section). The lineage commitment was evaluated by IF staining for the following germ layers markers: Sox17 for endoderm, Brachyury for mesoderm and Otx2 for ectoderm (Figure 2E; control negative staining of undifferentiated iPSCs is shown in Supplementary Figure S3). All tests confirmed the pluripotency, purity and stability of the five iPSC clones.

As newly established clones were validated as functional iPSCs, we utilized them to obtain muscle cells *in vitro*. Using a transgene-free method based on growth media, we differentiated iPSCs into muscle precursors (SCs), myoblasts and multinucleated myotubes (Figure 3A, 3B). In addition to the iPSCs prepared in this study, the previously published clones: E1M40 1.7, E1M40 1.9, E1M51 1.4, and E1M51 1.8 [25], which have the *EMD* mutation c.153del resulting in the absence of emerin, were also differentiated to muscle cells.

**Figure 3.**
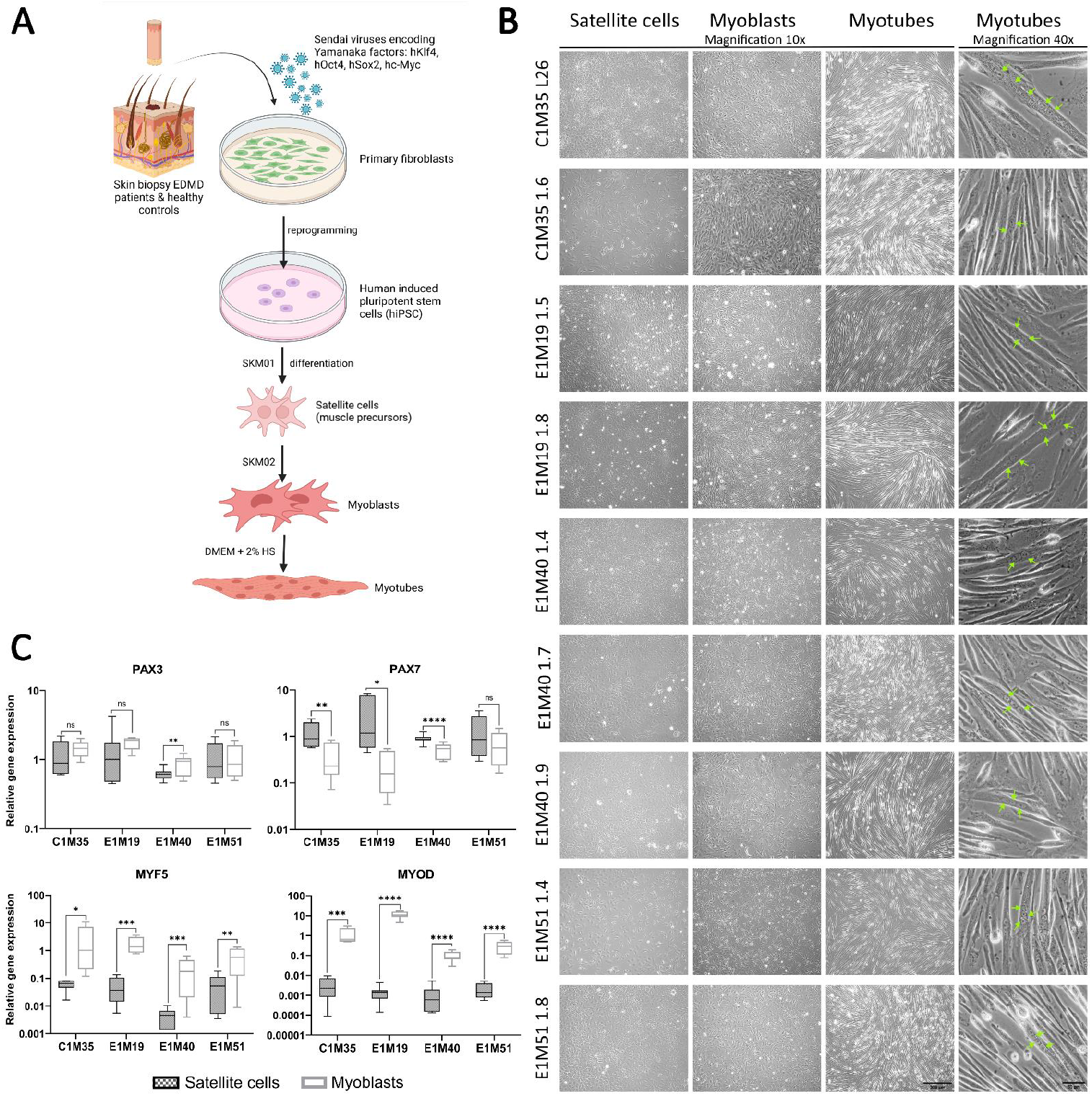
iPSCs differentiation into subsequent developmental stages of muscle cells. **(A)** Graphical representation of the reprogramming of fibroblasts from skin biopsies and differentiation process to subsequent developmental stages of muscle cells. **(B)** Images of SCs (day 6-7), myoblasts (day 6-7) and myotubes (up to 5 days in horse serum (HS)), transmission light, 10x magnification, the scale bar 300 μm, with multinucleated myotubes, 40x magnification, the scale bar 50 μm. Green arrows point to nuclei within an exemplary multinucleated myotube. **(C)** qPCR analyses of relative transcript levels of muscle early markers (Pax3, Pax7, Myf5, MyoD) in SCs (day 5-6) and myoblasts (day 6-7). Boxes depict 25th to 75th percentiles of relative gene expression with mean value marked with horizontal line, whiskers illustrate minimal and maximal values, asterisks indicate statistical significance ( * p ≤ 0.05, ** p ≤ 0.01, *** p ≤ 0.001, **** p ≤ 0.0001, ns – not significant). Samples were normalized to GAPDH and HPRT and referred to C1M35 donor cell lines; to SCs for Pax3 and Pax7 and to myoblasts for Myf5 and MyoD, n≥12 (≥4 biological replicates, 3 technical replicates).

Using qPCR, we confirmed the expression of muscle markers of particular stages of myogenesis. SCs express transcription factors Pax3 and Pax7 (Figure 3C), but not pluripotency marker Oct4 (analyzed by Western blotting (WB), Supplementary Figure S4). At the myoblast stage, the expression levels of myoblasts markers Myf5 and MyoD were significantly elevated in comparison to SCs (Figure 3C, at least 10-fold and 150-fold increase for Myf5 and MyoD, respectively). Pax3 expression level was sustained in myoblasts except E1M40, while Pax7 level decreased significantly (at least 2-fold) in all donors but E1M51. Pax3, Pax7 and Myf5 presence was also confirmed by WB (Supplementary Figure S4). For each clone, we obtained elongated myotubes (Figure 3B, right panel), Bbut the differentiation efficiency differed between samples. Depending on the clone, cells required 1-3 weeks of proliferation in myoblast medium, followed by differentiation, to be able to produce multinucleated myotubes. The prolonged culture in myoblast medium was necessary for E1M40 and E1M51 myoblasts as they did not differentiate into myotubes directly after reaching full confluency in myoblast medium, as C1M35 and E1M19 cells (Figure 4A).

**Figure 4.**
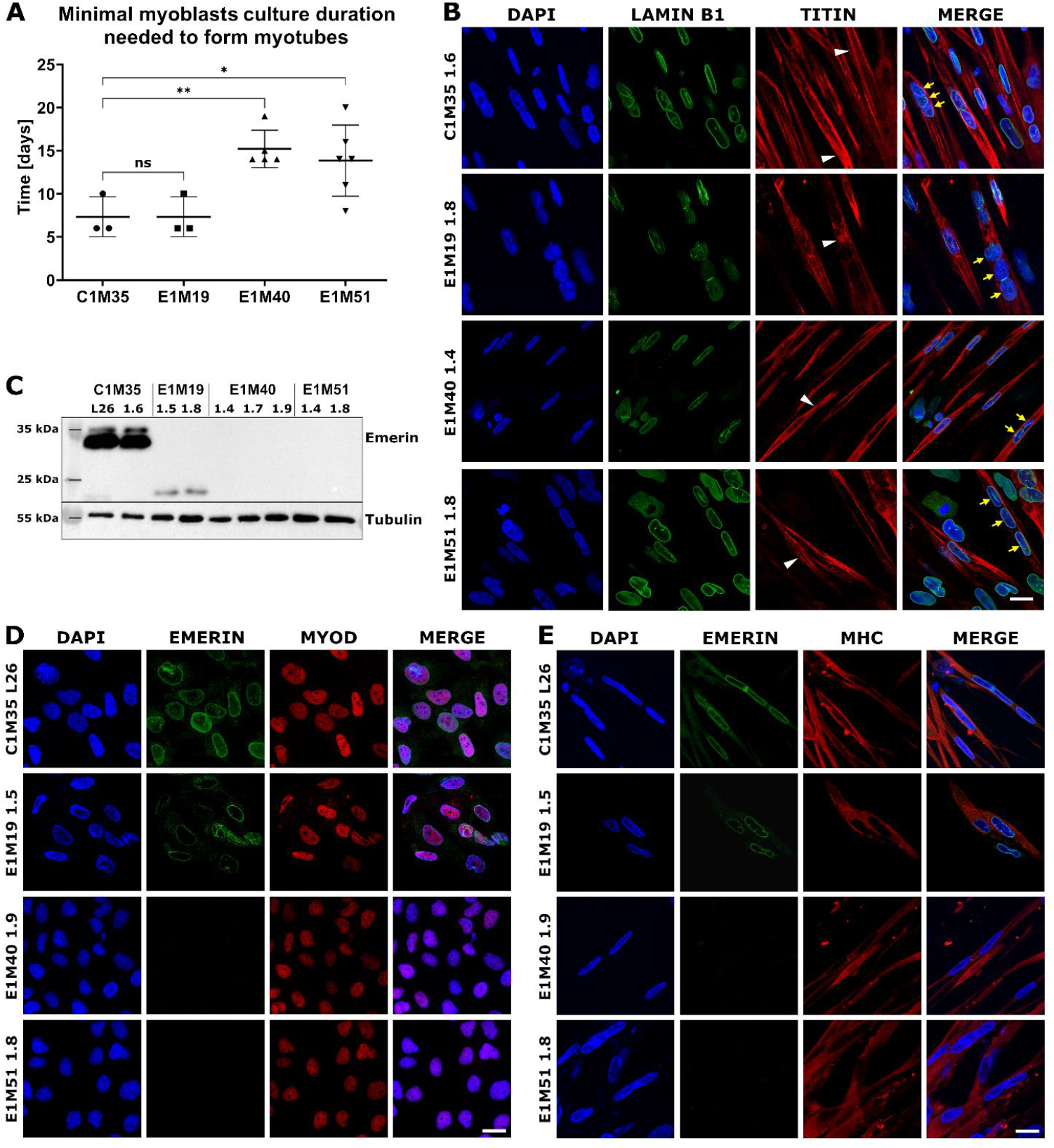
Characterization of myoblasts and myotubes obtained by iPSCs differentiation *in vitro*. Graph shows the minimal duration of myoblasts culture that is necessary for ability to form myotubes in subsequent differentiation medium. Each point indicates time spent by myoblasts in myoblast medium before performing successful differentiation to myotubes. The mean is marked with horizontal line, error bars show standard deviations, and asterisks indicate statistical significance (ns – not significant, * p ≤ 0.05, ** p ≤ 0.01); n≥3. **(B)** IF images show myotubes derived from patient’s cells, triple stained for chromatin (DAPI), lamin B1 (AF488) and titin (DyLight650). Titin staining shows sarcomere structure in myotubes (white arrowheads), while yellow arrows point to nuclei within an exemplary multinucleated myotube. The scale bar is 20 μm. **(C)** WB analysis of emerin level and size in myotubes. Full-length emerin was present in control cells, lower level of its truncated form was detected in E1M19 clones, while emerin was absent in E1M40 and E1M51 myotubes. IF images of myoblasts **(D)** and myotubes **(E)** triple stained for chromatin (DAPI), emerin (AF488) and muscle markers: MyoD (TRITC) for myoblasts and MHC (TRITC) for myotubes, confirming differentiation stage. Muscle cells show weaker signal for emerin in E1M19, and absence of signal in E1M40 and E1M51 clones. The scale bar is 20 μm.

Myotubes had an elongated morphology as expected, with the fraction of multinucleated cells (Figure 3B). IF analysis of myotubes revealed a sarcomere structure shown by titin bands in some cells (Figure 4B). Lamin B1 staining showed multiple nuclei in titin-positive cells, which confirms cells’ fusion. Using WB, we confirmed the presence of typical myotube markers: desmin, actinin, and MHC. All clones expressed lamins A/C, which is the main interaction partner of emerin, and its level increases during the differentiation (Supplementary Figure S5). In conclusion, we obtained mature myotubes from control and patients-derived cells.

Using WB and IF, we evaluated the localization and level of emerin in myoblasts and myotubes. In WB analysis (Figure 4C), the full-length emerin (254 amino acids, 29 kDa, migrating around 34 kDa due to posttranslational modifications) was detected only in control cells. For E1M19 cells we noted a lower level of a faster migrating form (less than 25 kDa) of truncated emerin, resulting from a frameshift from Glycine 151, followed by a STOP codon after 208 amino acids [32]. For clones E1M40 and E1M51, we did not observe any signal, suggesting a complete loss of expression of this protein as a result of mutation c.153del. We also analyzed emerin localization in myoblast and myotubes. In comparison to control cells C1M35, the intensity of the fluorescent signal for emerin was notably reduced in NE for E1M19 and completely disappeared in E1M40 and E1M51 cells (Figure 4D, E). All analyzed clones showed positive staining for the myoblast marker MyoD or myotubes marker MHC, respectively, confirming their muscle phenotype (Figure 4D, E).

## 3. Discussion

EDMD1, a rare genetic disorder belonging to laminopathies, still remains without a cure. Available treatments are limited to symptom-targeted therapies [33]. The precise molecular mechanisms underlying EDMD1 are not fully understood. The development of an appropriate research model is crucial for advancing our understanding of disease, which may also lead to the discovery of more effective therapies.

iPSC-based models have been established for a range of muscular dystrophies and laminopathies, including Limb-Girdle muscular dystrophy [34], Duchenne muscular dystrophy [35], Hutchinson-Gilford progeria syndrome or Dilated Cardiomyopathy [36], and also for EDMD type 2 (laminopathy caused by a mutation in *LMNA* gene encoding lamin A/C) [37, 38]. These models are widely used for *in vitro* differentiation into various tissues such as skeletal muscles, cardiomyocytes, adipocytes, and vascular smooth muscle cells, which serve as a model of tissues affected by a particular disease. Their utility in studying molecular mechanisms, testing candidate therapies, and exploring novel therapeutics, is invaluable [34, 36, 38]. However, an iPSC-based model for EDMD1 has not yet been developed.

Here, we present the first cellular model of EDMD1 based on iPSCs generated from skin fibroblasts obtained from EDMD1 patients, which was verified for proper differentiation to skeletal muscle cells. This model includes a collection of iPSC lines from three patients, two of whom are unrelated but share the same mutation. This is one of the most comprehensive iPSC-based models of laminopathies, ensuring replicates of clones and control cells reprogrammed simultaneously with the patient’s cells. All clones, including those described here for the first time and previously published by our group [25], have been fully characterized and validated, demonstrating the expected functional properties of iPSCs. Additionally, the cell collection consists of phenotypes of both emerin-null and truncated emerin, enabling the future studies of EDMD1 in the context of its genetic and phenotypic heterogeneity.

iPSCs may serve as a valuable starting point for various applications. In the case of EDMD1, tissues the most affected by the disease are skeletal muscles and the heart. In this work, we demonstrated the ability of iPSCs, which were generated by us, to differentiate through all stages of myogenesis, including SCs, myoblasts, and myotubes. This model allows for future in-depth analysis of myogenesis, which is difficult to achieve using other methods, particularly given the challenges associated with obtaining muscle biopsies from patients suffering from rare muscular dystrophies. The use of a transgene-free reprogramming approach minimizes potential artifacts from exogenous protein expression or genetic modifications, ensuring the disease mechanism remains unchanged.

An unexpected outcome obtained in our analysis was the sustained expression level of transcription factor Pax3 during the myoblast stage in comparison to SCs. Pax3 and Pax7 are typically considered as SCs markers, and their level usually decreases in myoblasts. However, their persistence in myoblasts has been also observed in other studies, e.g. in myoblasts and myotubes obtained by MyoD overexpression [19], or in fusing myoblasts [39]. In the model described here, the presence of Pax3 in myoblasts did not impair their differentiation ability to myotubes.

We observed that emerin-null myoblasts had delayed differentiation timing, as they needed twice longer exposure to myoblast medium before they could successfully differentiate into myotubes (Figure 4A). This suggests that our model reflects disease phenotypes. This variability in efficiency could be investigated further to determine the molecular mechanisms and link with altered emerin expression. In the previous studies utilizing different EDMD1 models, the alterations in multiple signaling pathways important for myogenesis and muscles regeneration, such as AKT, Wnt/β-catenin, MAPK/Erk, IGF-1, TGF-β, or Notch, were associated with mutations in *EMD* gene [5, 7, 40, 41]. Our model allows verifying these changes in pathways’ activity using cells with patient-specifics genetic background, at different stages of myogenesis. Such analysis may be a starting point for further explanation of the phenomenon of delayed differentiation observed by us.

In summary, to the best of our knowledge, we have established the first iPSC-based model of EDMD1, derived from three patients, that can successfully progress through multiple stages of myogenesis. These iPSCs also have the potential for differentiation into other tissues, including neurons, cardiomyocytes, and organoid skeletal muscle models. This new EDMD1 platform provides a valuable tool for advancing therapeutic development, including high-throughput pharmacological screening, as well as gene and cell therapy investigations.

## 4. Materials and Methods

### iPSCs generation and expansion

Patients’ cells (fibroblasts from skin biopsies) were obtained from Dr hab. Agnieszka Madej-Pilarczyk, Mossakowski Medical Research Institute, Polish Academy of Sciences, Warsaw, Poland. Fibroblasts were cultured in a fibroblast medium containing DMEM with high glucose 4.5 g/L (Gibco) supplemented with 10% fetal bovine serum (FBS, Gibco), 2 mM GlutaMAX (Gibco), penicillin (100 U/mL, Gibco) and streptomycin (100 μg/mL, Gibco). For reprogramming, fibroblasts were plated on 35 mm dishes for 2 days and then transduced with a set of non-integrating Sendai viruses (SeVs) using the CytoTune 2.0 Sendai reprogramming kit (Invitrogen) according to the manufacturer’s instructions. The medium was changed daily with the fresh fibroblast medium. On day 7, cells were plated on new dishes coated with Geltrex (Gibco). Starting the following day, the fibroblast medium was replaced with Essential 8 Medium (Gibco) every day. After 3-4 weeks individual iPSCs colonies were manually picked and transferred to new Geltrex-coated 24-well plates. iPSCs were expanded in Essential 8 Medium on Geltrex-coated dishes, they were dissociated with 0.5 mM EDTA in DPBS and split every 4-5 days at a ratio 1:8-1:12 using RevitaCell Supplement (Gibco). Fibroblasts and early passage iPSCs were cultured at 37°C in 5% CO_2_ and frozen at passage 4-5. Cells were expanded until at least passage 10 in 10% O_2_, 37°C, 5% CO_2_ and characterized between passages 13-20.

### In vitro differentiation to three germ layers

*In vitro* trilineage differentiation was carried out using the Human Pluripotent Stem Cell Functional Identification Kit (R&D Systems) according to the manufacturer’s instructions. Briefly, iPSCs were trypsinized and seeded on Geltrex-coated coverslips in a 24-well plate, at a density of approximately 9×10^4^ cells per well in Essential 8 medium with RevitaCell Supplement.

After obtaining 60-70% confluence for mesoderm and 70-80% for ectoderm and endoderm (24h/48h), E8 medium was replaced with proper differentiation medium (for endoderm, ectoderm or mesoderm, R&D Systems) and after 40 h for mesoderm and 72 h for ectoderm and endoderm coverslips were fixed with 4% PFA and stained for differentiation markers (as described in Immunofluorescence section).

### Differentiation of iPSCs into skeletal muscle cells

To generate skeletal muscle cells, the iPSC lines were seeded in 60 mm dishes coated with Geltrex, at a density of 2,500 or 5,000 cells per 1 cm^2^, depending on the clone, in Skeletal Muscle Induction Medium (SKM01, AMSBIO) supplemented with penicillin-streptomycin (Gibco). For the first 24 h, the RevitaCell Supplement was added to the medium. Then, the medium was changed every other day. Upon reaching full confluency (5-7 days), cells were collected using TrypLE Express Enzyme (Gibco) as SCs and seeded at a density of 2,500 cells per 1 cm^2^ of a 60 mm dish coated with Geltrex. Cells were seeded in the Myoblast Medium (SKM02, AMSBIO) supplemented with penicillin-streptomycin to induce differentiation into myoblasts. The medium was changed every 2-3 days. After the next 6-8 days, at the point of full confluency, the medium was changed to SKM03+ (AMSBIO) or DMEM-HS (DMEM high glucose (4.5 g/L, Gibco) supplemented with 2% HS (Gibco)), GlutaMAX Supplement and penicillin-streptomycin for myotube formation, and the cells were collected as myotubes after 3-7 days. Some clones needed additional culture time in SKM02 (up to 3 weeks), with passages every 3-5 days upon reaching full confluency, to be able to differentiate into myotubes. Cells were cultured in standard conditions (37°C, 5% CO_2_). Upon each differentiation stage cells were imaged with ZEN Software on ZEISS AxioVert Microscope with 10x objective and for myotubes additionally with 40x objective.

### Immunofluorescence (IF)

iPSCs were seeded on Geltrex-coated glass coverslips and cultured for 3-4 days until proper-sized colonies were formed. Myoblasts on Geltrex-coated glass coverslips in SKM02 medium were cultured 4-8 days to about 80% confluency when coverslips were fixed. To obtain myotubes specimens, myoblasts were seeded on Geltrex-coated glass coverslips and cultured for 3-4 days until reaching 100% confluency. Then the medium was changed to DMEM-HS or SKM03 and after 3-7 days myotubes were formed. Then coverslips were fixed in 4% paraformaldehyde for 20 minutes, washed with PBS, permeabilized with 0.5% Triton X-100 for 5 minutes at room temperature (RT) and washed again with PBS. Preparations were blocked for 30 minutes at RT with 1% donkey serum (DS, Gibco) or 1% FBS (Gibco) in PBS for germ layer markers and other stainings, respectively. Then preparations were incubated with primary unconjugated antibodies overnight at 4°C, washed with PBS and incubated with secondary antibodies or primary antibodies conjugated with a fluorophore for 1 h at RT, and washed again with PBS. Antibodies for iPSCs and germ layer markers were diluted in 1% DS in PBS (antibodies and dilutions used shown in Table 3). Coverslips were mounted on glass slides with the DABCO mounting medium (Fluka) with DAPI. Pluripotency markers were visualized with a Zeiss Axiovert A1 fluorescence microscope using a 10x objective, using ZEN Software. Myoblasts and myotubes, and germ layers staining was visualized with SP8 (Leica) or Stellaris (Leica) microscope using a 63x oil objective.

**Table 3.** Antibodies used for immunofluorescence (IF) and Western blotting (WB) staining.

|  | Antibody | Dilution | Company, cat# |
| --- | --- | --- | --- |
| Pluripotency markers | Rabbit anti Oct4<br>Mouse anti SSEA4 DyLight550 | 1:100 IF / 1:1000 WB<br>1:100 IF | Cell Signaling Technology cat# 2750<br>Invitrogen cat# MA1-021-D550 |
| iPSCs differentiation markers | Goat anti SOX17<br>Goat anti Otx2<br>Goat anti Brachyury | 1:10 IF<br>1:10 IF<br>1:10 IF | R&D Systems cat# AF1924<br>R&D Systems cat# AF1979<br>R&D Systems cat# AF2085 |
| Skeletal muscle markers and reference proteins | Mouse anti Pax3<br>Mouse anti Pax7<br>Mouse anti MyoD<br>Rabbit anti Myf5<br>Mouse anti MHC<br>Mouse anti desmin<br>Mouse anti actinin<br>Mouse anti myogenin<br>Rabbit anti GAPDH<br>Mouse anti actin α<br>Mouse anti tubulin α<br>Rabbit anti emerin<br>Rabbit anti lamin A/C<br>Rabbit anti lamin B1 | 1:1000 WB<br>1:200 WB (milk)<br>1:30 IF<br>1:2500 WB (milk)<br>1:10 IF / 1:500 WB<br>1:2500 WB<br>1:2000 WB (milk)<br>1:50 WB<br>1:10000 WB (milk)<br>1:1000 WB<br>1:1000 WB (milk)<br>1:400 IF / 1:1000 WB (milk)<br>1:4000 WB (milk)<br>1:100 IF / 1:500 WB | DSHB clone C2 [D1]<br>DSHB [D2]<br>Santa Cruz cat# sc-377460<br>Abcam cat# ab125078<br>DSHB clone MF-20 [D3]<br>Sigma Aldrich cat# SAB4200707<br>Sigma Aldrich cat# A7811<br>DSHB clone F5D [D4]<br>Sigma Aldrich cat# G9545<br>Sigma Aldrich cat# A2228<br>DSHB clone 12G10 [D5]<br>Cell Signaling Technology cat# 30853<br>Cell Signaling Technology cat# 2032S<br>Proteintech cat# 12987-1-AP |
| Secondary antibodies | Donkey anti-goat AF568<br>Donkey anti-rabbit TRITC<br>Goat anti-mouse IgM DyLight650<br>Donkey anti-rabbit AF488<br>Goat anti-rabbit HRP<br>Donkey anti mouse HRP | 1:200 IF<br>1:50 IF<br>1:50 IF<br>1:200 IF<br>1:10000 WB<br>1:10000 WB | Invitrogen cat# A11057<br>Jackson ImmunoResearch cat# 711-025-152<br>Invitrogen cat# SA5-10153<br>Jackson ImmunoResearch cat# 711-545-152<br>Jackson ImmunoResearch cat#111-035-144<br>Jackson ImmunoResearch cat# 715-035-151 |

Antibodies deposited to the Developmental Studies Hybridoma Bank (DSHB) by [D1] C.P. Ordahl, [D2] A. Kawakami, [D3] D.A. Fischman, [D4] W.E. Wright and J. Frankel / E.M. Nelsen. MHC - myosin heavy chain, HRP – horseradish peroxidase.

### Gene expression analysis

Total RNA was extracted from iPSCs using the Universal DNA/RNA/Protein Purification Kit (EurX). 1 μg of total RNA was reverse transcribed to cDNA with Maxima First Strand cDNA Synthesis Kit for RT-qPCR (Thermo Scientific). Real time qPCR was performed using SYBR Green Master Mix (Applied Biosystems) and Quant Studio 5 Real-Time PCR System (Applied Biosystems); all samples were analyzed in four technical replicates. The reaction parameters were as follows: 50°C for 2 minutes, 95°C for 2 minutes, followed by 40 cycles of denaturation at 95°C for 15 seconds and annealing/extension at 64°C for 30 seconds. Additionally, the 3-step melt curve was performed (95°C for 15 seconds, 60°C for 1 minute, 95°C for 15 seconds). Pluripotency markers’ relative gene expression was normalized to Glyceraldehyde-3-Phosphate Dehydrogenase (*GAPDH*) and referred to an established iPSCs line E1M40 1.7 (UWRBTi003-B [25]). The sequences of the designed primers are listed in Table 4.

**Table 4.**
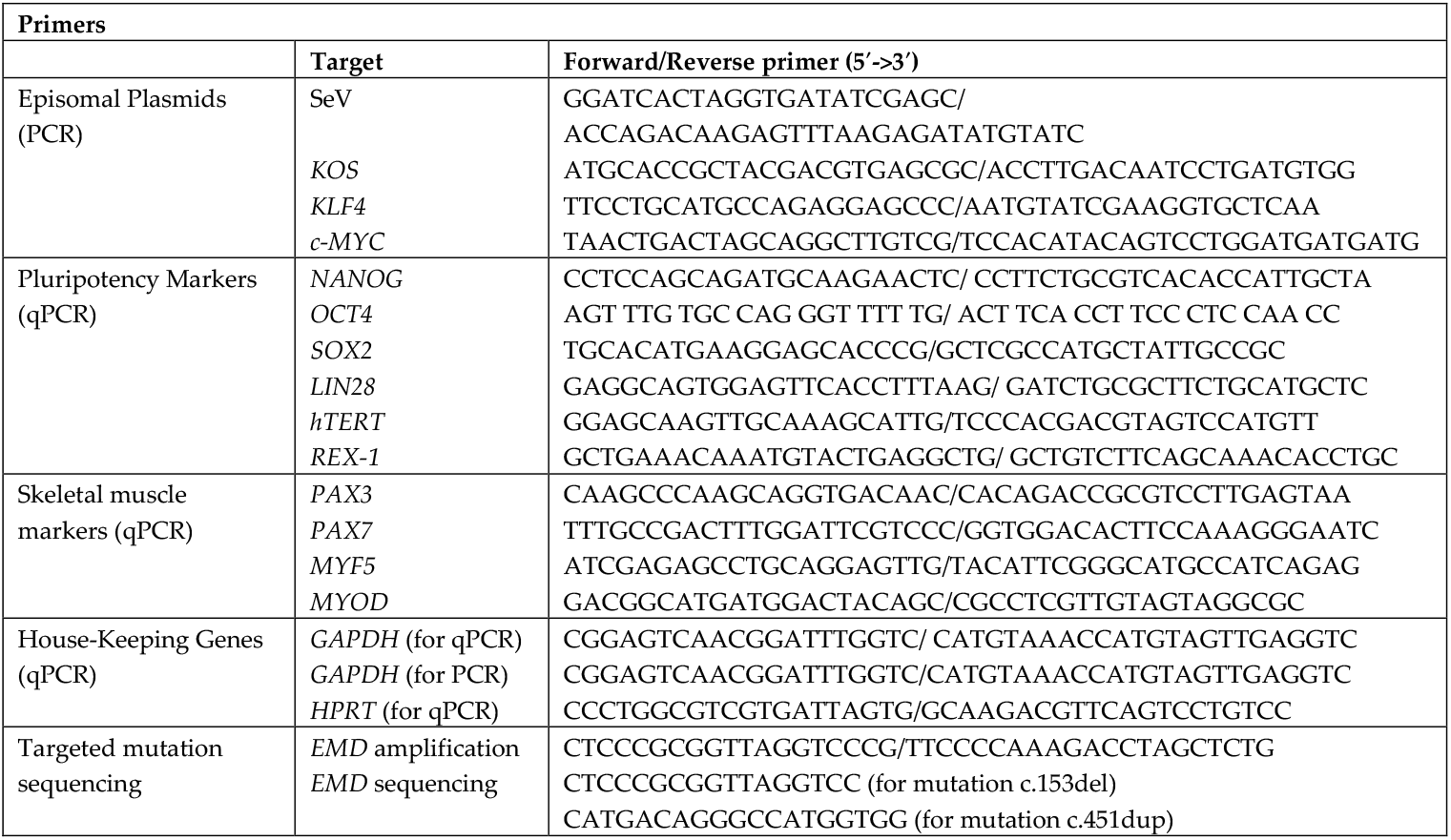
Primers. List of all primes used for qPCR, *EMD* sequencing and episomal vector presence detection. SeV sequence is a genomic sequence of Sendai virus. KOS sequence contains human KLF4, OCT3/4, and SOX2.

For the analysis of muscle markers, the total RNA was extracted using Direct-zol RNA Microprep Kit (Zymo Research). 500 ng of the isolated RNA was reverse transcribed as described above. qPCR was performed as described with samples being analyzed in at least 4 biological replicates, each in three technical repeats. Relative gene expression was normalized to GAPDH and Hypoxanthine phosphoribosyltransferase (HPRT) and referred to C1M35 donor cell lines (to C1M35 SCs for Pax3 and Pax7 and to C1M35 myoblasts for Myf5 and MyoD). Primers’ sequences are presented in Table 4. Statistical significance was calculated using multiple t-test in GraphPad Prism (version 8.4.3).

### Short tandem repeats (STR) analysis

The profiling of the human cell lines was performed by Microsynth and Eurofins companies using highly polymorphic STR loci. They were amplified using the PowerPlex 16 HS System (Promega). Fragment analysis was done on an ABI3730xl (Life Technologies) and the resulting data were analyzed with GeneMarker HID software (Softgenetics).

Eurofins performs genotyping according to ANSI/ATCC standard ASN-0002 testing 16 DNA markers using the Applied Biosystems AmpFLSTR Identifiler Plus PCR Amplification Kit system.

### Sequencing

Genomic DNA was isolated using Universal DNA/RNA/Protein Purification Kit (EurX), then *EMD* gene was amplified with Phusion HSII DNA Polymerase (Thermo Scientific), with GC buffer and 3% DMSO and previously published [16] primers listed in Table 4. Reactions were performed using thermocycler TOne 96G (Biometra) with the following parameters: 98°C for 5 minutes, followed by 35 cycles of denaturing at 98°C for 15 s, annealing at 62°C for 30 s, and extension at 72°C for 90 s and 10 minutes final extension at 72°C. PCR product was purified with PCR/DNA Clean-Up Purification Kit (EurX) and sequenced by Microsynth company with sequencing primers shown in Table4.

### Karyotyping

iPSC lines and fibroblasts were treated with 0.67 μg/mL colcemid (BioWest) for 2.5 hours, then dissociated by Trypsin-EDTA (Gibco) and centrifuged at 1400 rpm, 7 minutes. After incubation in hypotonic solution (Ohnuki’s solution) and fixation in methanol:glacial acetic acid (3:1) mixture (Chempur), microscopic slides were prepared. G-banded metaphase analysis was performed according to the International System for Human Cytogenomic Nomenclature 2020 (ISCN 2020; AGT manual [42], [43]) employing the Imager.M1 (Zeiss) microscope and the Ikaros software (version 6.3, Metasystems DE). Up to 30 metaphases from iPSCs (resolution 400-450 bands) and at least 15 metaphases from fibroblasts (resolution 350-400) were karyotyped by an experienced cytogeneticist.

### The episomal vectors’ presence

Episomal vectors presence was analyzed by PCR performed for iPSCs cDNA using 5 pairs of primers shown in Table 4 four pairs recognizing reprogramming vectors (sequences from CytoTune-iPS 2.0 Sendai Reprogramming Kit) and 1 pair for *GAPDH* as control. Reactions were performed with Taq polymerase (EURx), with 35 cycles of amplification, using thermocycler TOne 96G (Biometra) with the following parameters: 95°C for 5 minutes, followed by 35 cycles of denaturing at 95°C for 30 s, annealing at 55°C (*c-MYC, GAPDH, SeV, KOS*) or 63°C (*KLF4*) for 30 s, and extension at 72°C for 30 s and 5 minutes final extension at 72°C. Normal human dermal fibroblasts (Lonza) were used as a negative control, cells collected shortly after SeV transduction were used as a positive control.

### Mycoplasma test

Mycoplasma PCR test was performed on culture supernatants (at least 48h without medium change and confluence at least 80%) using Kit Mycoplasma PCR Detection Kit (Abcam) according to the manufacturer’s instructions.

### Western blotting (WB)

Cell pellets were resuspended in hot Laemmli lysis buffer (50 mM Tris-HCl pH 6.8; 10% glycerol; 2% SDS). Lysates were boiled for 5 minutes (96°C) and cooled. BCA assay (Thermo Scientific) was performed for 10x diluted samples, according to the manufacturer’s instructions, in a 96-well plate format. After measurements, samples were supplemented with DTT to a final concentration of 50 mM and boiled again. 25 μg of protein extracts were separated via SDS-PAGE on 6%-12% gels, transferred to nitrocellulose membranes (0.45 μm pore sizes), and blocked in 5% non-fat milk in PBS with 0.075% Tween 20 (PBST) for 1 h, RT. Primary antibodies were applied overnight at 4°C in PBST (or in 5% non-fat milk as marked in the Table 3), and secondary antibodies conjugated with HRP were applied for 1 h, RT in 5% milk in PBST buffer (antibodies and dilutions shown in Table 3). The proteins were visualized using ECL substrate (Bio-Rad) and analyzed with ImageLab (Bio-Rad).

## Supplementary Materials

**Supplementary Table S1**. STR profiles summary of characterized iPSC clones and their parental fibroblasts.

**Supplementary Figure S1**. Negative control of staining performed for pluripotency markers (Oct4 and SSEA4) on HeLa cells.

**Supplementary Figure S2**. The analysis of parental fibroblasts.

**Supplementary Figure S3**. Negative control of staining of three germ layers markers (Sox17, Brachyury and Otx2) performed for undifferentiated IPC clone E1M40 1.4.

**Supplementary Figure S4**. The analysis of protein levels in iPSCs-derived SCs and myoblasts.

**Supplementary Figure S5**. WB analysis of myogenesis markers levels in myotubes differentiated from IPSC.

## Authors’ contribution

KP, MM and RR designed the study. CB and MM generated iPSCs. KP, ML and MR expanded iPSCs. ML, MR, IŁ and AS performed iPSCs validation tests. AS, KP, ML, MM differentiated iPSCs to muscle cells. AS, KP, JH, AK and ML performed muscle cells analysis. VD was responsible for the resources. KP supervised the study. KP and MM wrote the original draft of the manuscript. All authors reviewed the manuscript. KP, MM and RR provided funding.

## Funding

The study was funded mainly by the National Science Centre grant 2021/43/D/NZ5/01634 SONATA "Identification of molecular mechanisms underlying the phenotype development of Emery-Dreifuss muscular dystrophy” granted to KP.

Research project was partly supported by the program "Excellence initiative – research university” for years 2020-2026 for University of Wrocław granted to KP, by the grant KNOW for the National Leading Scientific Center for Biotechnology from the Polish Ministry of Science granted to MM and RR, the Statutory grant for research from the Polish Ministry of Science, and the Polish National Science Center MINIATURA 2 granted to MM.

## Acknowledgments

KP is thankful to Professor Francesco Saverio Tedesco and his team for sharing their expertise on iPSCs culturing and muscle cells differentiation *in vitro*. Authors are thankful to Dr hab. Agnieszka Madej-Pilarczyk for providing patients’ cells.

## Conflicts of Interest

The authors declare no conflict of interest.

## Disclaimer/Publisher’s Note

The statements, opinions and data contained in all publications are solely those of the individual author(s) and contributor(s) and not of MDPI and/or the editor(s). MDPI and/or the editor(s) disclaim responsibility for any injury to people or property resulting from any ideas, methods, instructions or products referred to in the content.

## Notes

### Competing Interest Statement

The authors have declared no competing interest.

### Summary of Updates

The revised version was broadly corrected.

