## Supplementary data for "Human iPSC-derived muscle cells as a new model for investigation of EDMD1 pathogenesis"

| LOCI | E1M40 FIBRO | E1M40 1.4 | C1M35 FIBRO | C1M35 1.6 | C1M35 L26 | E1M19 FIBRO | E1M19 1.5 | E1M19 1.8 |
|---|---|---|---|---|---|---|---|---|
| D8S1179 | 13 15 | 13 15 | 12 | 12 | 12 | 12 13 | 12 13 | 12 13 |
| D21S11 | 28 31 | 28 31 | 30 32.2 | 30 32.2 | 30 32.2 | 29 31 | 29 31 | 29 31 |
| D7S820 | 9 | 9 | 11 12 | 11 12 | 11 12 | 11 12 | 11 12 | 11 12 |
| CSF1PO | 11 12 | 11 12 | 11 13 | 11 13 | 11 13 | 10 11 | 10 11 | 10 11 |
| D3S1358 | 18 20 | 18 20 | 16 19 | 16 19 | 16 19 | 14 15 | 14 15 | 14 15 |
| TH01 | 7 9 | 7 9 | 7 8 | 7 8 | 7 8 | 6 9 | 6 9 | 6 9 |
| D13S317 | 12 13 | 12 13 | 8 12 | 8 12 | 8 12 | 11 12 | 11 12 | 11 12 |
| D16S539 | 12 | 12 | 11 | 11 | 11 | 9 11 | 9 11 | 9 11 |
| vWA | 14 15 | 14 15 | 16 18 | 16 18 | 16 18 | 16 16 | 16 16 | 16 16 |
| TPOX | 10 11 | 10 11 | 8 | 8 | 8 | 8 11 | 8 11 | 8 11 |
| D18S51 | 11 15 | 11 15 | 12 16 | 12 16 | 12 16 | 13 14 | 13 14 | 13 14 |
| AMEL | X Y | X Y | X Y | X Y | X Y | X Y | X Y | X Y |
| D5S818 | 12 | 12 | 11 14 | 11 14 | 11 14 | 11 13 | 11 13 | 11 13 |
| FGA | 21 23 | 21 23 | 22 23 | 22 23 | 22 23 | 20 23 | 20 23 | 20 23 |

**Table S1**. STR profiles summary of characterized iPSC clones and their parental fibroblasts.

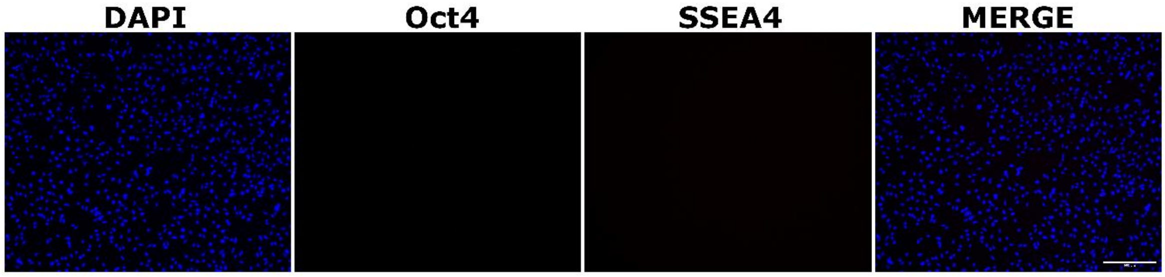


**Figure S1.** Negative control of staining performed for pluripotency markers (Oct4 and SSEA4) on HeLa cells. This result confirms specificity of staining in Figure 1B. Scale bar is 500 µm.

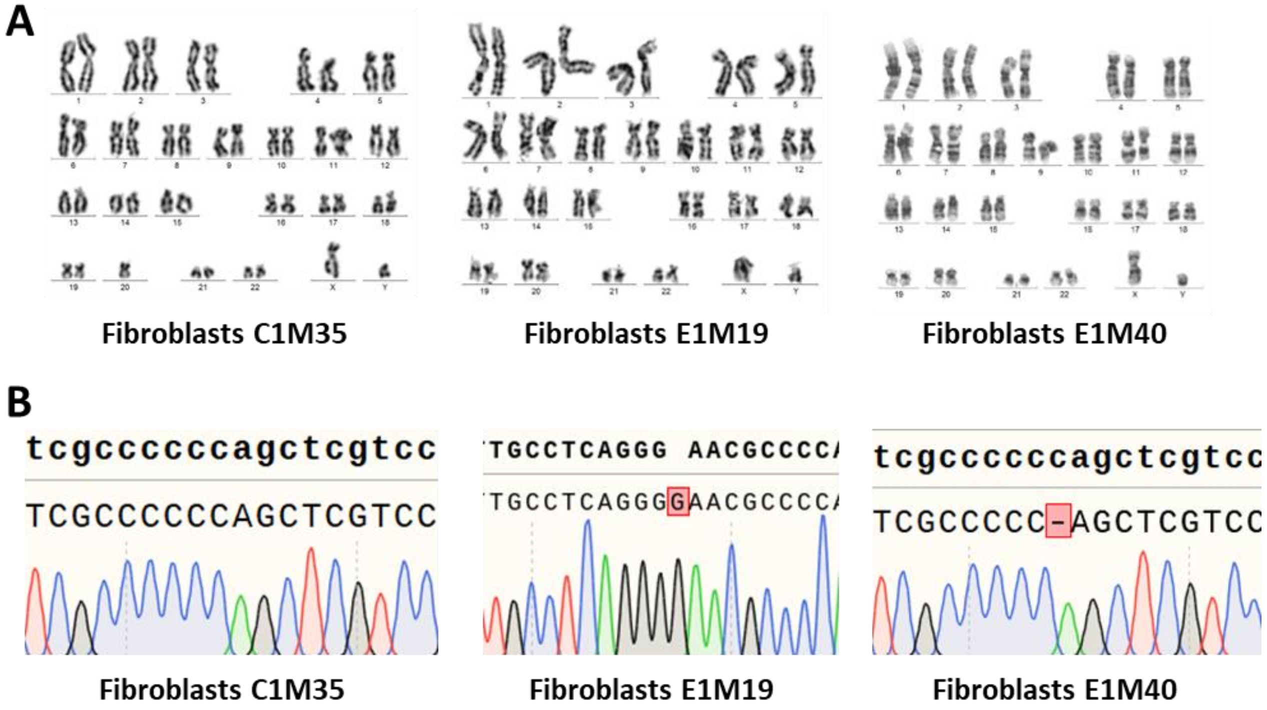


**Figure S2**. Analysis of parental fibroblast lines. (A) Karyotypes of parental fibroblasts. (B) DNA sequencing results of *EMD* gene in parental fibroblasts. The upper sequence shows the reference wild-type *EMD*, while the bottom sequence shows the result of DNA sequencing.

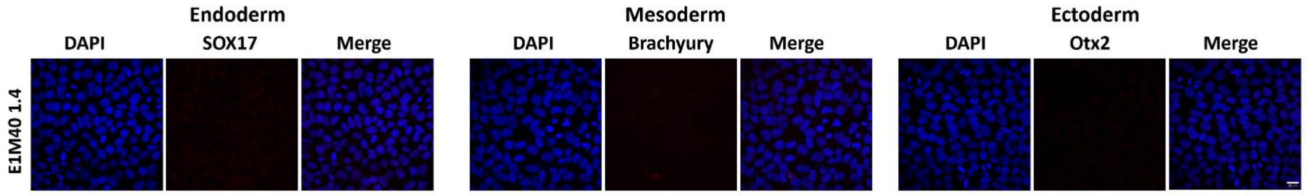


**Figure S3.** Negative control of staining of three germ layers markers (Sox17, Brachyury and Otx2) performed for undifferentiated iPSC clone E1M40 1.4. This result confirms specificity of staining in Figure 2E. Scale bar is 20 μm.

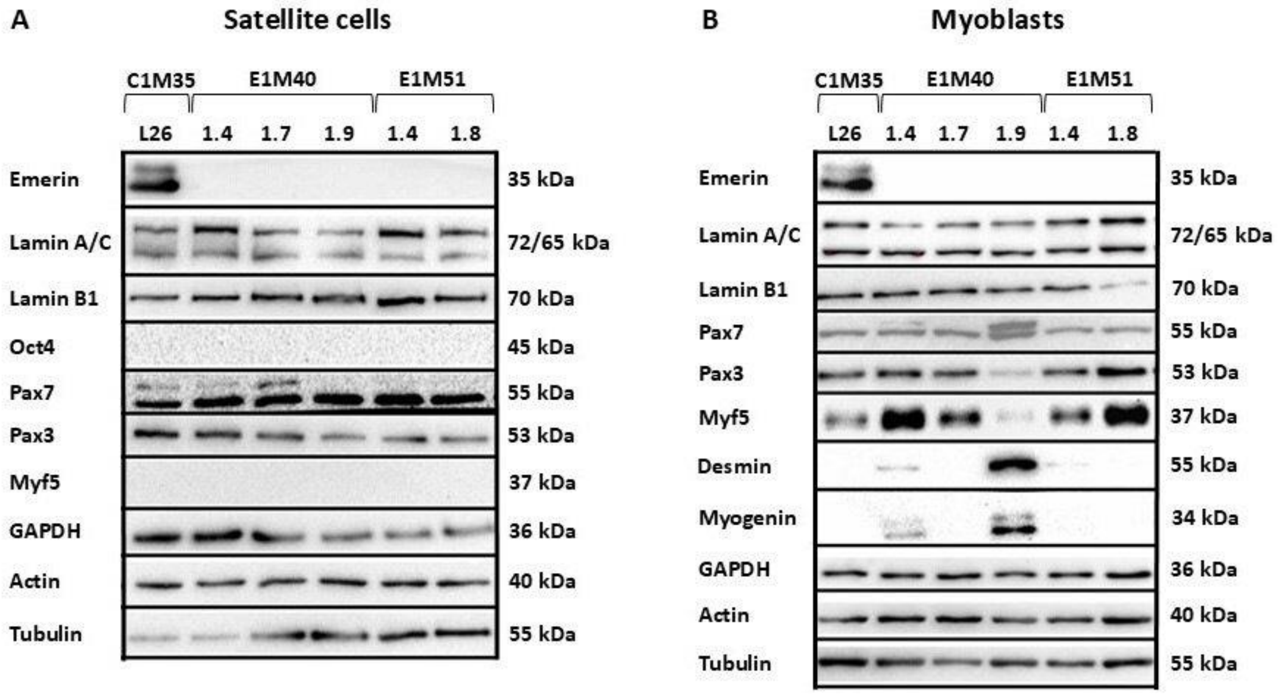


**Figure S4.** The analysis of protein levels in iPSC-derived SCs and myoblasts. One clone of iPSCs from a healthy donor (C1M35 L26) and five clones from two patients (E1M40 1.4, 1.7 and 1.9, and E1M51 1.4 and 1.8) with mutation c.153del (emerin null) were differentiated to SCs (A) and myoblasts (B). The samples were collected at different time points, once the cells reached 100% confluency, which occurred within 5-8 days for satellite cells and 4 to 6 days for myoblasts. Then, the expression of the following proteins was determined using western blot: nuclear-envelope proteins emerin, lamin A/C and lamin B; iPSCs marker Oct4; SCs markers Pax3 and Pax7; myoblasts marker Myf5; myotubes markers desmin and myogenin; and controls GAPDH, actin and tubulin. In both SCs and myoblasts, emerin is present only in the control line, while, as expected, lacking in the patient's cells (any faster-migrating form was not detected in optimized conditions; data not shown). In SCs, the expression of markers of particular stage of differentiation is proper - lack of iPSCs marker Oct4, expression of progenitor cells markers Pax3 and Pax7, and lack of Myf5 - marker of myoblasts - the subsequent stage of myogenesis. In myoblasts, Pax3 and Pax7 are still present, and the Myf5 is already expressed. In some clones, markers of myotubes (a subsequent stage of differentiation) - desmin and myogenin are present, which indicates that some cells started spontaneous differentiation.

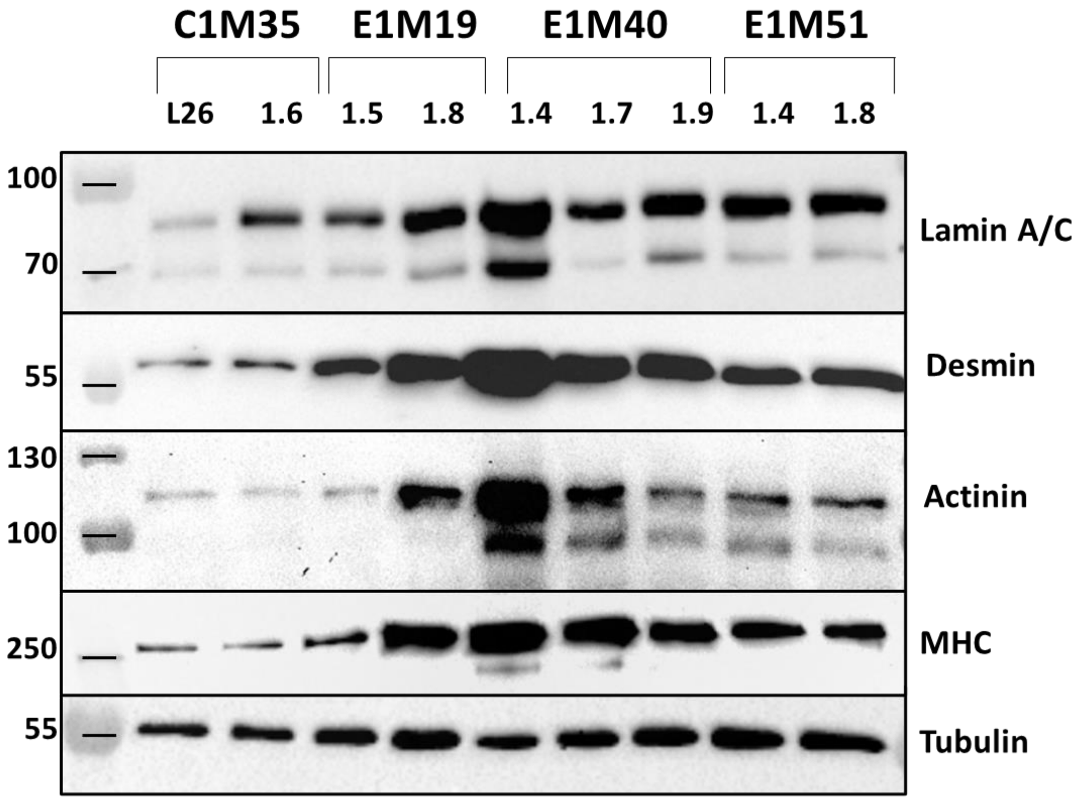


**Figure S5.** WB analysis of myogenesis markers in myotubes differentiated from iPSCs.
WB analysis was performed for two clones from healthy donor (C1M35) and seven clones from three patients. (E1M19, E1M40, E1M51). The presence of lamin A/C, desmin, MHC and actinin were confirmed in all clones, while proteins' levels depend on clone properties and differentiation time. Tubulin was used as loading control (the same staining as in the Figure 4C). Samples corresponding to WB presented on Figure 4C.
iPSC-derived myoblasts were cultured in myoblast medium for 1-3 weeks, depending on the clone, and differentiated to myotubes. C1M35 and E1M19 clones were able to differentiate into myotubes directly after medium change from myoblast medium to myotube medium, while E1M40 and E1M51 clones needed additional passaging and culturing in myoblast medium. WB present comparison of myotube samples obtained in the first possible time point for each clone.
